# Distinct patterns of thought mediate the link between brain functional connectome and psychological well-being

**DOI:** 10.1101/762344

**Authors:** Deniz Vatansever, Theodoros Karapanagiotidis, Daniel S. Margulies, Elizabeth Jefferies, Jonathan Smallwood

**Affiliations:** Institute of Science and Technology for Brain-inspired Intelligence, Fudan University, Shanghai, PR China, 200433; Department of Psychology, University of York, York, United Kingdom, YO10 5DD; Institut de Cerveau et de la Moelle pinire, Centre National de la Recherche Scientifique, Paris, France, 75013

**Keywords:** functional connectivity, functional magnetic resonance imaging, graph theory, mental health, resting state, patterns of thought

## Abstract

Ongoing thought patterns constitute important aspects of both healthy and abnormal human cognition. However, the neural mechanisms behind these daily experiences and their contribution to well-being remain a matter of debate. Here, using resting state fMRI and retrospective thought sampling in a large neurotypical cohort (n = 211) we identified two distinct patterns of thought, broadly describing the participants current concerns and future plans, that significantly explained variability in the individual functional connectomes. Consistent with the view that ongoing thoughts are an emergent property of multiple neural systems, network-based analysis highlighted the central importance of both unimodal and transmodal cortices in the generation of these experiences. Importantly, while state-dependent current concerns predicted better psychological health, mediating the effect of functional connectomes; trait-level future plans were related to better social health, yet with no mediatory influence. Collectively, we show that ongoing thoughts can influence the link between brain physiology and well-being.

## INTRODUCTION

Recent advances in functional magnetic resonance imaging (fMRI) methods and data analysis techniques have facilitated a new era in the characterisation of the neural representations that underlie human cognition and behaviour [1]. Big data-driven approaches are utilised to explore whole-brain functional interactions during unconstrained states of rest, explaining considerable levels of population-wise variation in complex traits, including intelligence [2], personality [3], daily habits [4] and self-perceived quality of life [5]. In addition to helping researchers derive novel theories on healthy brain processing, one important goal of these functional connectomic mapping initiatives is to devise neural, cognitive and behavioural links to better understand mental health disorders, and to identify “at-risk” groups for future preventative measures [6, 7]. However, most studies often neglect the fact that periods of unconstrained states of rest can be characterised by patterns of self-generated thoughts that are also associated with well-being, which may have unique neurocognitive correlates.

Research from psychology suggest that the ability to self-generate patterns of cognition is a core element of our mental lives, occupying a considerable portion of our daily mentation [8–10]. Critically, these thoughts are particularly prevalent in situations with low external demands, such as periods of wakeful rest [11] i.e. the conditions when resting state functional data is most commonly recorded. Importantly, measures of such experiences have a wide range of momentary correlates including indicators of stress [12], ongoing physiology [13] and task performance [14], and also have documented links to both beneficial and deleterious psychological traits. For example, patterns of ongoing thought have been previously linked to individuals’ ability to plan for future goals [15], wait for long-term rewards [16], and devise creative solutions to both personal [17] and social problems [18]. Other forms of self-generated mentation are associated with unhappiness [19], as well as poor performance in sustained attention tasks [20] or measures of fluid intelligence [21]. In fact, disruptive thought patterns are reported to underlie the absent-minded mistakes in our everyday functioning [22, 23], including traffic accidents [24] or medical malpractice [25], and to form a potential basis for cognitive impairments reported in attention-deficit/hyperactivity disorder [26, 27] and clinical depression [28].

Furthermore, there is now emerging evidence that link patterns of neural activity to trait variance in specific patterns of thoughts. These investigations highlight that patterns of ongoing thought have complex and often heterogeneous neural correlates [29, 30], which depend on the functional interaction of multiple neural and cognitive components [31–33]. For example, Wang and colleagues identified distinct patterns of ongoing thoughts that were linked to reductions in both within and between network connectivity for neural systems important for external attention (ventral, dorsal and fronto parietal systems), which was in turn related to reduced performance in tasks of cognitive aptitude [34]. Other studies have illustrated that different types of spontaneous thought depend on differential patterns of functional connectivity between regions important for memory (e.g. temporal lobe) and those that form the core associative cortices [30, 35, 36]. Notably, tasks such as creative idea generation [37] and future planning [38], that are collectively associated with unconstrained thought [17, 39], depend on similar patterns of interaction between regions of the default mode with regions of the fronto-parietal network.

Converging body of evidence from contemporary cognitive neuroscience, therefore, highlight that (*a*) patterns of neural activity at rest have associated patterns of thought, and (*b*) that both neural and self-report descriptions of patterns of unconstrained activity are predictive of a wide range of psychological features of an individual, including mental well-being. Collectively, these observations highlight the need to understand the extent to which relationships between neural activity at rest and aspects of psychological traits are dependent upon the nature of patterns of ongoing experience that emerges while neural activity is recorded. Our current study reflects an attempt to address this issue by quantifying the relationship between patterns of thoughts during rest, and the associated ongoing neural activity, and then exploring whether descriptions of neural organisation gained in this fashion is associated with variation in measures of psychological trait, in this case self-perceived mental well-being.

In the analysis of a large neurotypical sample (n = 211) we found differential profiles of functional interactions amongst multiple neural systems that explained individual variation amongst two distinct patterns of thought, broadly corresponding to the participants current concerns and future plans. Using a test-retest sample (n = 40) we found reasonable concordance between the tendency to generate future plans at rest and their brain basis, indicating that these measures are likely to reflect a trait. On the other hand, current concerns were not stable across individuals, but showed evidence of common changes in terms of the pattern of thought and the pattern of functional connectivity, suggesting that these neurocognitive links reflect a more transient state. Finally, we found that while current concerns indirectly mediated the effect of brain functional interactions on psychological well-being, future plans showed no such mediation effect.

## RESULTS

### Dimensions of variation in ongoing thought patterns

With the aim of investigating the differential influence of distinct thought patterns on the link between brain functional network topology and mental well-being, we collected 9-minutes of fMRI data from a large cohort of neurotypical participants (n = 211) during a period of wakeful rest. This neural measure was complemented with a session of retrospective thought sampling, administered immediately after the resting state scanning, as well as self-assessed ratings of psychological and social well-being on a cross-culturally validated questionnaire from the World Health Organization Quality of Life (WHOQOL) group [40], collected outside the scanner in a separate behavioural session. The workflow for the data collection and analysis techniques utilized in this study is presented in Figure 1 with further details provided in the Methods and Supplementary Information. Parts of this dataset have been previously employed in our prior investigations [27, 32, 34, 41].

**FIG 1.**
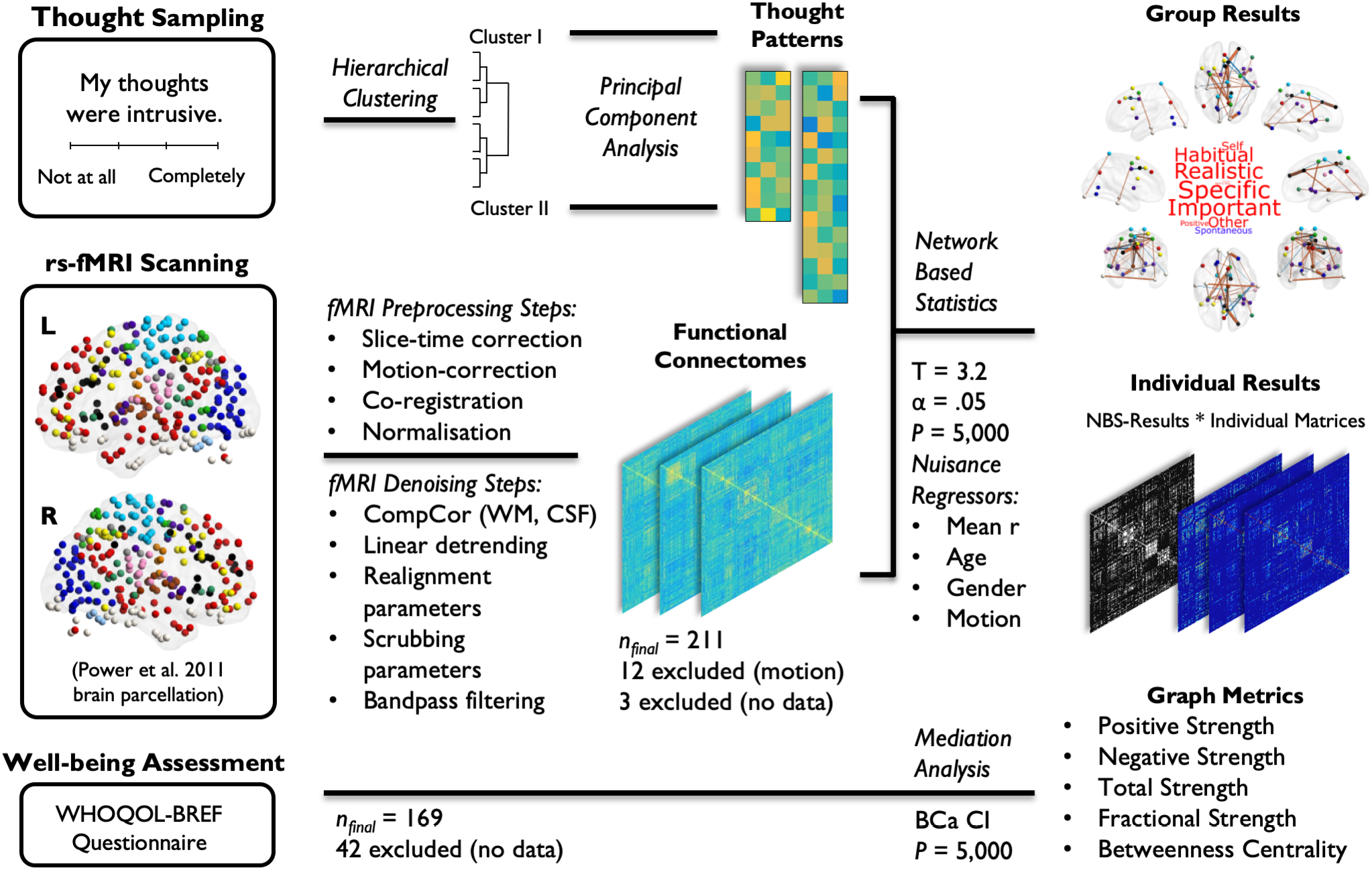
Experimental data collection and analysis pipeline. A large-cohort of participants (n = 211) were fMRI scanned during a state of wakeful rest for a total of 9-minutes. A comprehensive pipeline of fMRI preprocessing and data denoising procedures were followed in order to ensure maximal removal of nuisance variables. The thought sampling scores collected at the end of the resting state scanning session were first hierarchically clustered into two major groups, each of which were then reduced to three patterns of thought using principal component analysis (PCA). Fully-connected, weighted correlation matrices based on the Power et al. 2011 parcellation (264 regions of interest) scheme and the individual component scores on the identified patterns of thought were then carried forward on to Network-Based Statistics (NBS) with the aim of identifying components of brain connections that related to the participants thought patterns. T-tests were carried out for all six patterns of thought with an initial T-score of 3.2, and a significance level of p < .05 over 5,000 permutations (mean connectivity, age, gender and percentage of invalid scans based on the composite motion score from the scrubbing procedure were entered as group-level nuisance regressors). The identified brain components that significantly related to the participants thoughts were first characterized at the group-level and then used as mask graphs to create thresholded connectivity matrices for each participant. Network metrics of positive, negative, total and fractional strength (i.e. the ratio of positive to negative strength) as well as betweenness centrality were measured on individual thresholded matrices and were further used to characterize the identified neurocognitive profiles. Linear regressions and mediation analyses were then employed to investigate the mediatory effect of thought patterns on the link between brain connectivity and psychological and social well-being as measured using a cross-culturally validated World Health Organization Quality of Life group (WHOQOL-BREF) questionnaire.

Our first objective was to establish the dimensions of variation within the participants ongoing thought patterns. The employed retrospective thought sampling method required participants to subjectively characterize their thoughts using 4-choice (Likert scale) ratings on a set of questions derived from our previous investigations [18, 30, 33, 39, 42] (Supplementary Table S1). With the aim of reducing dimensionality and improving interpretability, these ratings were first hierarchically clustered and then decomposed into distinct dimensions using principal component analysis (PCA), which established the main patterns of thought reported in our sample (Fig. 2). The total number of thought patterns were selected using scree-plots, based on the eigenvalue (>1) and the explanatory power gained by each additional decomposition (Supplementary Fig. S1-2). Typical ratings from participants who scored highest on a given thought pattern are provided in Supplementary Figure S3.

**FIG 2.**
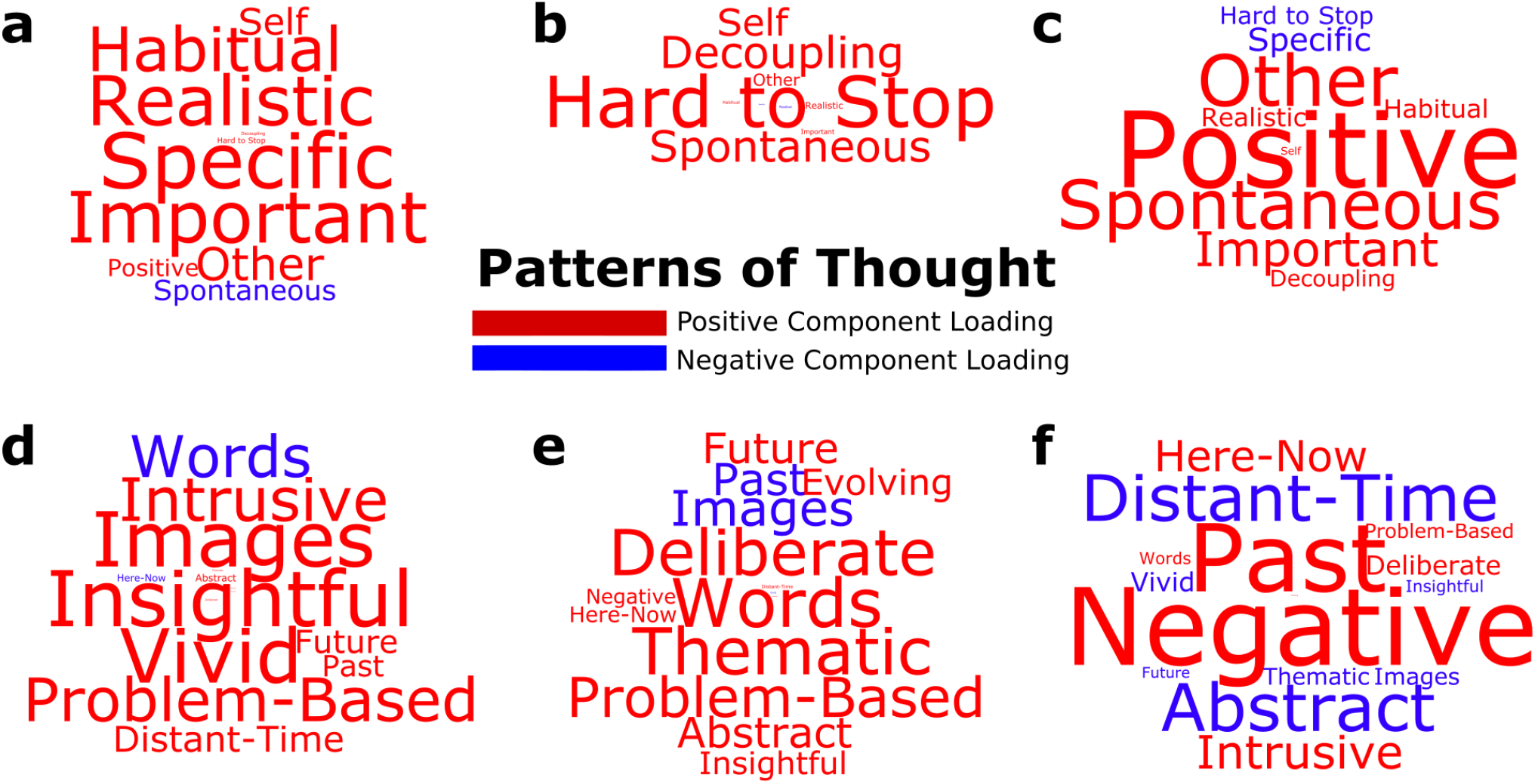
Decomposing distinct patterns of thought. Following the initial hierarchical clustering of the participants ratings on the experience sampling questionnaire, a total of six patterns of thought were identified using PCA. Three principle components in each of the two clusters explained 51% and 35% of the variability in the data, respectively. The Varimax rotated component loadings are visualized using word clouds. While the size of the text refers to the relative strength of the component loadings, positive and negative loadings are indicated via red and blue fonts, respectively. The components high-lighted (*a*) important/specific thoughts, (*b*) perceptually-decoupled/hard-to-stop thoughts, (*c*) positive/spontaneous thoughts, (*d*) insightful/image-based thoughts, (*e*) deliberate/verbal thoughts, and (*f*) negative/past-related thoughts. The individual variation on these patterns of thought were carried forward on to the NBS analysis as between-subject explanatory variables.

Principal Components Analysis (PCA), applied separately to each of the two initial hierarchical clusters on the self-reported ratings, revealed three patterns of thought each (a total of six), explaining 51% and 35% of the variance, respectively. The identified patterns highlighted aspects of ongoing thoughts that are commonly characterized in the existing literature [43]. These encompassed thoughts that were important and specific (Fig. 2a), perceptually decoupled and hard-to-stop thoughts about the self (Fig. 2b), positive, spontaneous thoughts about others (Fig. 2c), insightful and image-based thoughts (Fig. 2d), deliberate, verbal thoughts about the future (Fig. 2e), and negative past-related thoughts (Fig. 2f).

### Distinct patterns of thought link to differential functional connectomic profiles

Next, we determined the extent to which the human functional connectomes were associated with individual variation on the identified patterns of thought. We used a whole-brain parcellation scheme [44]. Defining each of the 264 brain regions as nodes, and Pearson correlation coefficients amongst them as edges, we constructed fully-connected, weighted functional connectomes for each participant. Utilizing Network-Based Statistics (NBS) [45], we entered the individual component scores for all six patterns of thought as variables of interest in a regression model, while removing the effects of nuisance variables including mean connectivity, age, gender and the composite motion score (i.e. percentage of invalid volumes identified in the motion artifact detection procedure) (Fig. S4). This revealed two distinct patterns of thought that were significantly related to differential profiles of brain functional connectomic organisation (T_threshold_ = 3.2, 5,000 permutations, α = .05). Although we report on the initial T_threshold_ = 3.2 threshold, comparable results for T_threshold_ = 3.1 and T_threshold_ = 3.3 are presented in the Supplementary Results section (Fig. S5).

The first thought pattern that was significantly associated with neural patterns reflected high loadings on “habitual”, “realistic”, “specific” and “important” elements of ongoing cognition, broadly corresponding to the construct of current concerns that has been argued to play an important motivational role in the content and form of ongoing thoughts [46]. Population variation in this experience was linked to a combination of positive and negative connections from a wide range of both unimodal and transmodal brain regions (Fig. 3a) (T_threshold_ = 3.2, 5,000 permutations, p = .015). These included areas associated with visual, auditory, somato-motor, dorsal/ventral attention, cingulo-opercular, salience, fronto-parietal, default mode, subcortical networks and a set of regions with memory-related functions, with the strongest link observed between the right middle frontal and middle cingulate gyri (salience network).

**FIG 3.**
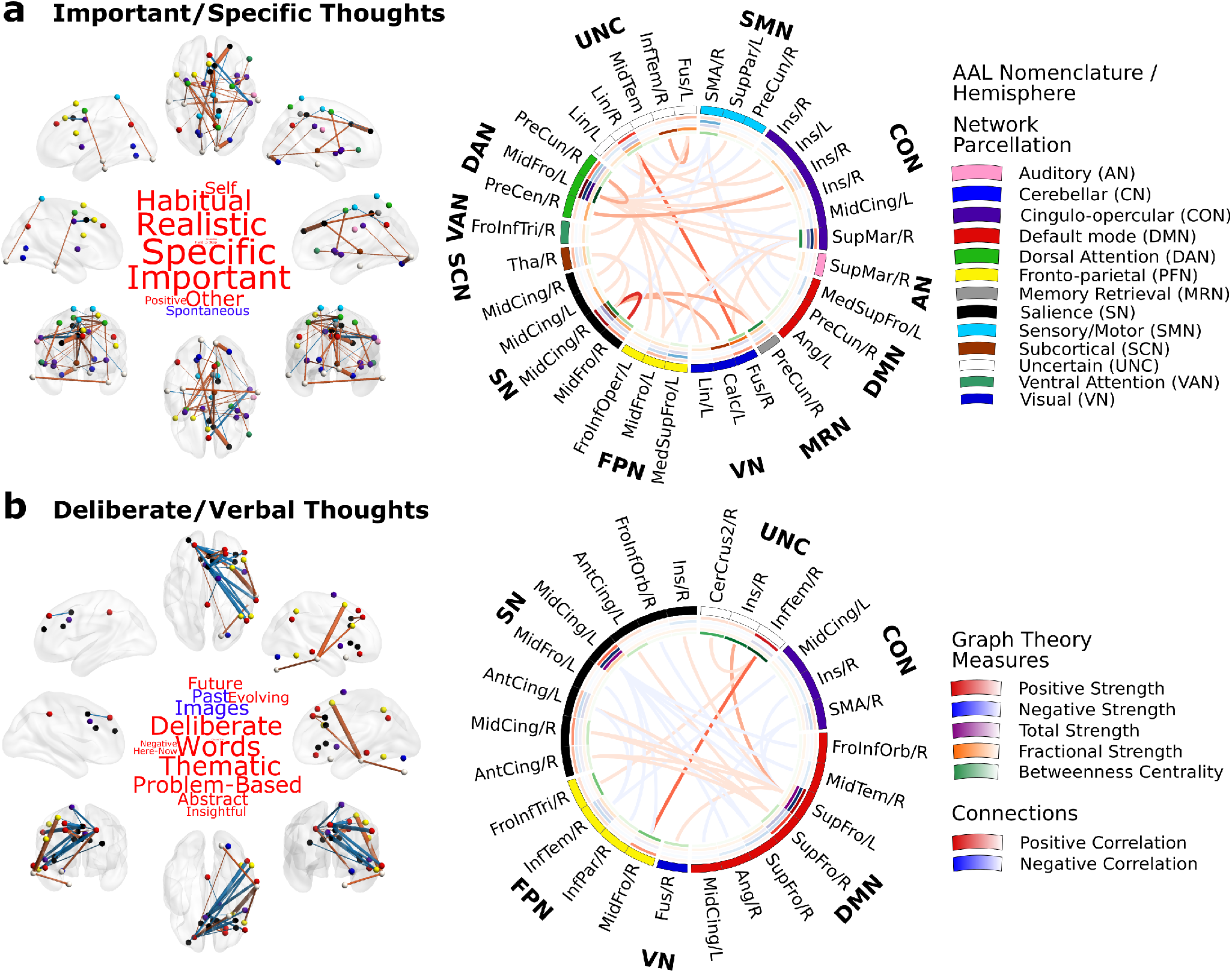
Distinct patterns of thought are linked to differential profiles of functional connectomic organisation. NBS identified two connected components of brain functional interactions at rest that significantly related to the participants scores on distinct patterns of thought, which highlighted (*a*) important, specific, realistic and habitual thoughts about the self and others that were not spontaneous; as well as (*b*) deliberate, verbal, thematic, abstract, problem-based, and future-oriented thoughts in words that were not about the past or in images. The average connectivity patterns of these brain graphs are visualized on an MNI152 smoothed glass brain, with the nodes color-coded according to the original Power et al. 2011 parcellation scheme, and the positive/negative connections coloured in red and blue, respectively. The average connectivity patterns of these brain graphs and the average graph theory metrics calculated across participants are visualized in circular representations. The Automated Anatomical Labeling (AAL) nomenclature, original network parcellation, average positive (red), negative (blue), total (purple) and fractional strengths (orange) as well as betweenness centrality (green) of each region in this brain component are listed around the rings. The average positive and negative connections between these regions are represented by links colour coded red and blue respectively.

Overall, this neural component was composed of more positive than negative connections and involved a set of distributed brain regions that encompassed a diverse set of large-scale brain networks. Characterization of this brain component using network neuroscience measures revealed that an area in the left middle frontal gyrus, belonging to the dorsal attention network, showed the highest positive, negative, and total strength as well as betweenness centrality. This highlights the left middle frontal gyrus as a key node in the functional connectivity patterns linked to ongoing thoughts about the individuals’ current concerns. In addition, the highest fractional strength (i.e. the ratio of positive to negative strength) was observed in a region belonging to the salience network, namely the right middle cingulate gyrus.

The second pattern of thought that related to neural function, reflected high loadings on the “future” rather than the “past”, as well as “deliberate” and “verbal”focus on “problems”, which is broadly consistent with studies of ongoing thought that emphasize a deliberate focus on future plans [15, 47]. This thought pattern was related to the functional interaction of brain regions from a small set of large-scale brain networks mainly belonging to the higher-order transmodal cortices (T_threshold_ = 3.2, 5,000 permutations, p = .026) (Fig. 3b). Both positive and negative links between regions belonging to the default mode, fronto-parietal, salience, cingulo-opercular and visual regions correlated with higher scores reported on this pattern of thought, with the strongest link observed between regions in the right inferior temporal and middle frontal gyri. Relative to the “current concerns” component, this brain component was composed of more negative connections and was drawn from a more localized set of brain regions from a smaller number of large-scale brain networks. Network-level analysis indicated that a left superior frontal gyrus region belonging to the default mode network showed the highest positive, negative and total strength, while a left middle cingulate region (salience network) displayed the greatest fractional strength. Betweenness centrality, however, was highest on a right inferior temporal gyrus region.

### Test-retest reliability of thought patterns and functional connectomic organisation

Next, we examined the stability of the two neurocognitive measures over time by exploring their intraclass correlation across two sessions in 40 participants for whom a second assessment was performed (i.e. resting state scan and thought sampling). For “important and specific thoughts about the self and others” (i.e. current concerns), individual variability on this thought pattern displayed no significant intraclass correlation (ICC = .14, CI [-.18, .43], p = .20), however, the associated brain connectivity (natural log of the fractional strength) was correlated between the first and second assessment sessions (ICC = .60, CI [.36, .76], p < .0001). Importantly, there was a significant positive correlation between the change in component scores on this thought pattern and the change in brain connectivity between the two assessment sessions (Pearson r = .40, p = .011) (Fig. 4a). Together this pattern is broadly consistent with a more transient state.

**FIG 4.**
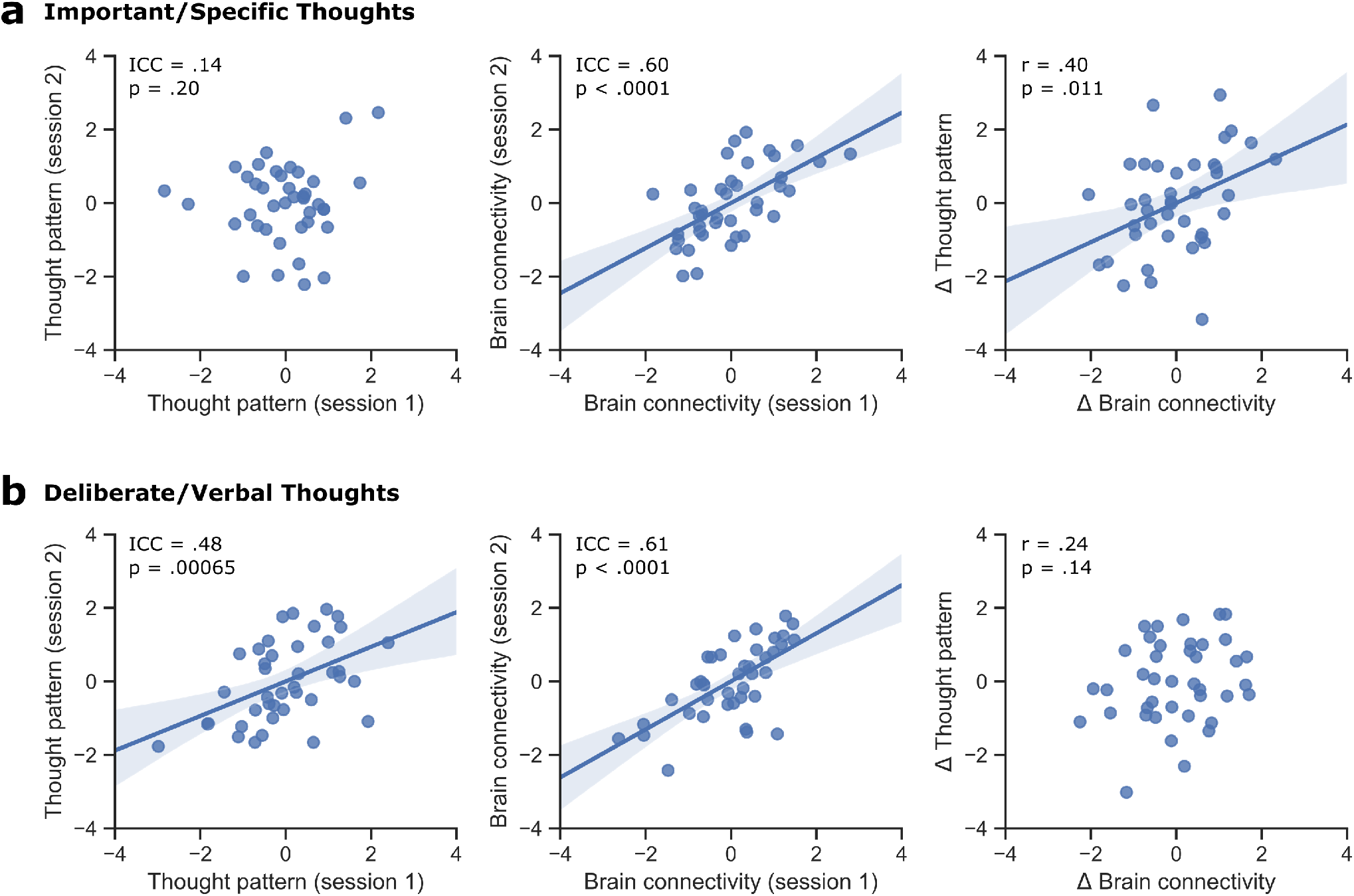
Test-retest reliability of the identified thought patterns and the associated functional connectomic organisation. A second session of resting state fMRI scanning and thought sampling was carried out for a total of 40 participants. (*a*) For important/specific thoughts, while individual component scores on this thought pattern did not show a significant intraclass correlation (ICC = .14, CI [-.18, .43], p = .20), the associated brain connectivity pattern (natural log of fractional strength) was largely consistent between the two repeated assessment sessions (ICC = .60, CI [.36, .76], p < .0001). Moreover, the change in thought pattern was positively related to the change in brain connectivity (Pearson r = .40, p = .011). (*b*) For deliberate/verbal thoughts, both the component scores on this thought pattern (ICC = .48, CI [.21, .69], p = .00065) and the associated brain connectivity (ICC = .61, CI [.38, .55] p < .0001) showed significant intraclass correlations. However, there was no significant link between the change in this thought pattern and the change in brain connectivity between the two assessment sessions (Pearson r = .24, p = .14).

For “deliberate verbal thoughts about the future” (i.e. future plans) on the other hand, individual variability on both the thought pattern (ICC = .48, CI [.21, .69], p = .00065) and the associated brain connectivity (ICC = .61, CI [.38, .55] p < .0001) were consistent across the two assessment sessions. However, there was no significant correlation between the change in component scores on this thought pattern and the change in brain connectivity (Pearson r = .24, p = .14) (Fig. 4b). Collectively, these analyses show that the patterns of thought captured by “deliberate and verbal thoughts about the future” and their neural representations show greater traitlike stability over time than the participants state-like “important specific thoughts about the self and others”.

### Distinct patterns of thought mediate the effect of brain connectivity on well-being

Finally, having identified two patterns of ongoing thought, each with an associated profile of complex brain functional interactions at rest, we tested whether these neurocognitive metrics derived from our study had mediatory influences on measures of mental health and well-being in daily life, as indicated by a cross-culturally validated World Health Organization questionnaire (i.e. WHOQOL-BREF). Incorporating the relative importance of both positive and negative connections [48, 49], we first assessed the predictive power of fractional strength on psychological and social well-being via linear regressions, followed by mediation analyses to examine the indirect influence of brain connectivity on well-being, mediated by the participants reported patterns of thought.

For “important and specific thoughts about the self and others” (i.e. *current concerns*), the fractional strength (natural log) of the associated brain connectivity component significantly predicted the participants’ psychological well-being score β = .52, r = .19, p = .014), while no significant link was observed with social well-being β = .075, r = .030, p = .699). A mediation analysis indicated that there was a significant indirect effect of brain connectivity on psychological well-being, mediated by the individuals’ scores on the identified pattern of thought β′ = .35, 95% CI [.029, .68]) (Fig. 5a). For “deliberate and verbal thoughts about the future” (i.e. *future plans*) on the other hand, the fractional strength (natural log) of the identified component of brain connectivity significantly predicted the participants social well-being score on the WHOQOL-BREF β = .38, r = .17, p = .024), while no significant link was observed with psychological well-being β = -.069, r = .028, p = .718). Moreover, a mediation analysis revealed no evidence for an indirect effect of brain connectivity on social well-being through the individuals’ scores on the identified thought pattern β’ = .11, 95% CI [-.071, .32]) (Fig. 5b).

**FIG 5.**
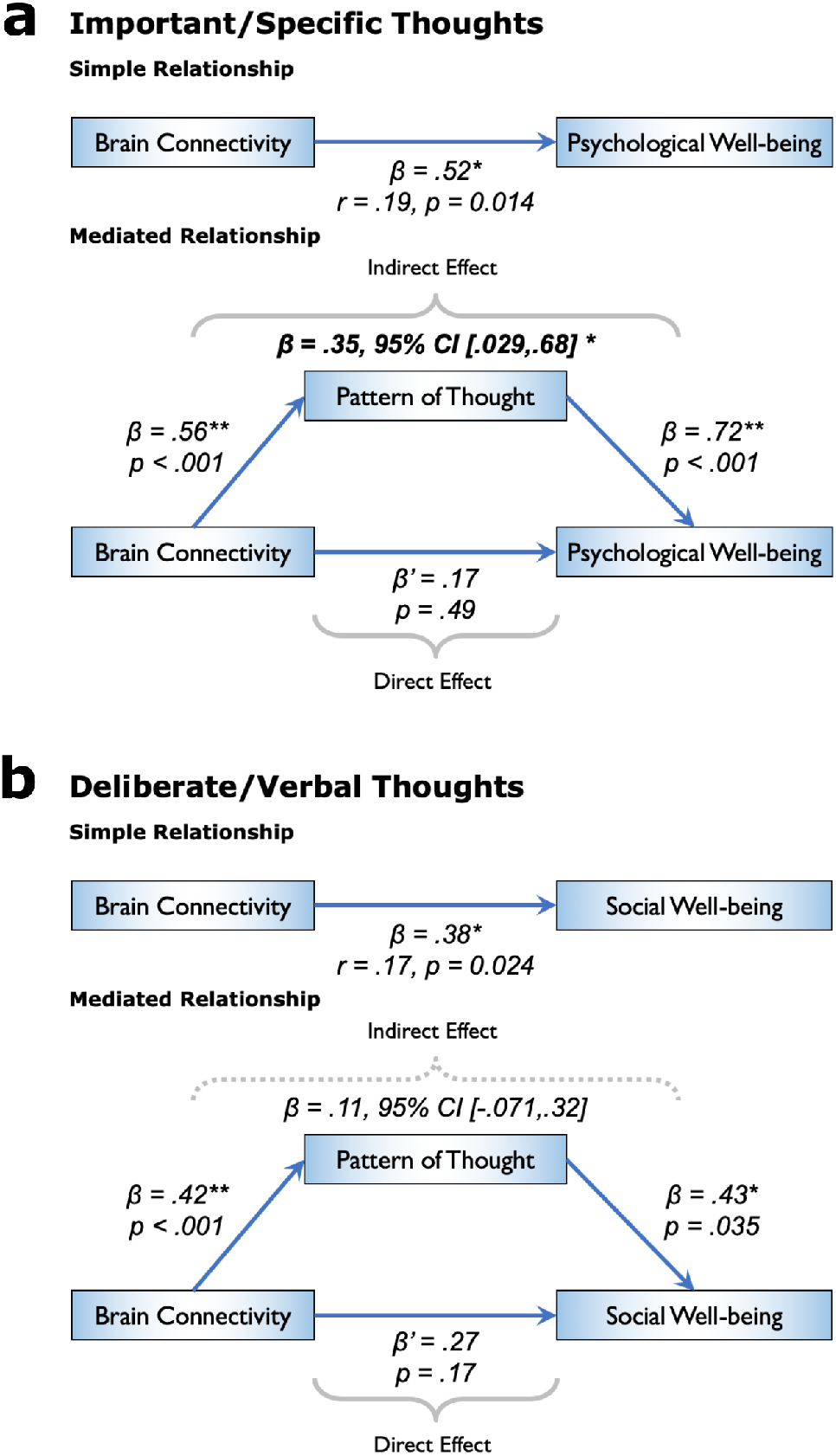
Model of the functional connectome as a predictor of psychological and social well-being, mediated by patterns of thought. While brain connectivity refers to the natural log of the fractional strength, pattern of thought represents the individual scores on the identified thought pattern, and the psychological/social well-being measures are self-reported ratings on the WHOQOL-BREF questionnaire. Only a subset of the participant cohort who fully completed the well-being questionnaire (n = 169) was utilised in this analysis. The calculation of confidence intervals (CI) for the indirect effect was based on the percentile bootstrap estimation approach with 5,000 samples. (*a*) A significant indirect effect of brain connectivity on psychological well-being was observed, mediated through the participants important/specific thoughts, broadly related to their current concerns. (*b*) There was no significant indirect effect of brain connectivity on social well-being, mediated through the participants deliberate/verbal thoughts on their future plans.

## DISCUSSION

The aim of this study was to examine whether accounting for the patterns of ongoing thoughts that individuals experience during periods of wakeful rest (e.g. resting state fMRI scanning) could allow for a more nuanced understanding of the relationships between aspects of psychological functioning, patterns of neural organisation and mental health. To achieve this goal, we first identified patterns of neural connectivity at rest that varied with aspects of self-reported experience during this period. One dimension of variation was along patterns of thinking that was indicative of a focus on current concerns, which was dominated by connections from the middle frontal gyrus, a region within the salience network. This neural pattern was a stable feature of the assessed individuals, while the pattern of thinking did not depict such reliability. Nevertheless, both neural and self-report patterns changed together across time, indicating that this neurocognitive profile described a transient mapping between brain and experience. A second pattern, associated with deliberate thoughts about the future, was dominated by functional connections from the superior frontal gyrus, situated within the default mode network. Both neural and experiential features of this mode were consistent across individuals and showed little evidence of common changes over time, suggesting a neural pattern that was a relatively stable trait. Importantly, these neural components had differential associations with measures of mental health and well-being. The transient neurocognitive component, linked to the participants focus on current concerns, was significantly associated with aspects of self-reported psychological well-being. Mediation analysis indicated that this brain-behaviour relationship was fully mediated by the associated descriptions of ongoing experience. In contrast, while the dimensions of variation linked to deliberate problem solving was associated with social well-being, this relationship was independent of the associated patterns of experience.

Together these analyses indicate that important aspects of the commonly reported relationships between brain and behaviour can be understood through their associations with patterns of ongoing thoughts that participants experience during fMRI scanning. First, our study shows that neural profiles, identified through their association with self-reported experience, have differential associations with aspects of well-being. Self-reports are obviously subject to factors that impact upon their credibility, however, neural patterns activity identified in this fashion have the advantage that they are embedded in a cognitive context without the need for reverse inference [50]. Thus, experience sampling provides a complimentary method for determining whether the source of observed neural patterns are cognitive in nature, or emerge for other reasons, such as cardiovascular function or motion related confounds [51]. Second, our study demonstrates that experience sampling is sensitive to patterns of neural activity that are stable across time and others that are transient. Our approach, therefore, may have direct relevance to studies that aim to explore the long-term stability of patterns of functional organisation, or those that explore long-term changes in neural function. Based on the current data, for example, experience sampling may provide a reasonably direct way to test hypotheses into why certain neural patterns vary in their consistency across time. Despite the inherent weakness associated with retrospective experience sampling [52], our study shows that the information it provides can address important shortcomings of the conceptual interpretations placed on functional connectivity patterns derived from resting state analysis. Given the negligible cost associated with acquiring descriptions of experience after resting state scans, and their prevalence as a tool of cognitive neuroscience, we see no reason why this method should not be employed as a community standard in similar studies moving forward.

As well as highlighting the value of experience sampling to studies investigating the relationship between functional organisation and behaviour, our study provides valuable information into the neural processes that contribute to different types of self-generated thought. Current concerns, either in the form of unfulfilled goals or personally relevant information, occupy a significant portion of the thoughts we experience in our daily lives, potentially constituting a determining factor in the functional outcomes of our ongoing cognition [28, 46]. In line with these results, a key dimension of thought pattern that was reported by the participants in our cohort was related to “important and specific thoughts about the self and others” or more generally their current concerns. Our results revealed that important functional connections was observed in the salience network, commonly implicated in the detection of behaviourally important, internal or external stimuli for the coordination of neural resources [53]. This neural system has been shown to causally influence the functional interaction between default mode and fronto-parietal networks that are commonly anti-correlated at rest [54]. We recently combined momentary experience sampling with online neural activity and found that a pre-frontal region of this network was associated with the ability to prioritise patterns of episodic social thoughts during periods when external demands were reduced [55]. Together with such evidence our study suggests that the role of the saliency network in patterns of ongoing thought emerges from its general capacity for prioritizing patterns sensory, memorial and affective content [52] that is motivated by their contextual relevance [56, 57].

A second component of “deliberate and verbal thoughts about the future” was linked to the connectivity of a limited number of nodes in transmodal cortices including regions of the default mode, cingulo-opercular, salience and fronto-parietal networks. This neural pattern was dominated by connections from a region of the superior frontal gyrus, within the default mode network. A deliberate focus on future goals reflects our ability to simulate and envision the consequences of our actions based on our prior experiences [58]. Importantly, similar interactions between the default mode network and systems linked to executive control (e.g. fronto-parietal and salience networks) are observed when participants engage in tasks that mimic this type of thought, such as creative problem-solving [37], visuospatial [59] as well as autobiographical planning [38], and imagining reward outcomes [60]. This pattern of thought was predictive of better social well-being, an observation consistent with studies linking patterns of ongoing thought to social problem solving [18], and our ability to infer the actions and mental states of others [61]. This component was also characterised by both positive as well as negative brain connections. Negative or anti-correlations have been historically assumed to arise from the analysis techniques employed, and head-motion that is thought to lead to spurious connectivity measures [62]. However, recent reports suggest a neurophysiological basis and a potential cognitive importance of such anti-correlations in healthy brain processing [48, 49, 63], and our results raise the possibility that the tuning of interactions between neural systems may give rise to different types of ongoing experience a hypothesis which requires further investigation. Furthermore, recent perspectives on the generation and maintenance of thought patterns highlight the vital importance of considering the dynamic nature of ongoing cognition and the within individual variation in this process [52]. Although retrospective thought sampling methods provide the advantage of acquiring measures related to an undisrupted period of unconstrained cognition, online thought sampling and the assessment of the link between neural and experiential dynamics might constitute a fruitful route in deciphering the within and between-individual variability in ongoing cognition and the underlying neural mechanisms [64].

In summary, we have shown that distinct patterns of thought are reflected in the underlying brain functional connectomes at rest and that certain types of these experiences may mediate the influence of intrinsic brain connectivity organization on well-being. Our results highlight the importance of considering thought patterns when establishing predictive relationships between functional connectomes and complex traits, but also suggest that taking this aspect of human cognition into consideration may lead to better characterization of neural finger prints of the connectome, with the potential for more useful clinical markers. In the future, it may be possible to tailor self-reported questions based on their neural associations, allowing development of measures targeting particular psychiatric populations with well-established neural hypotheses, such as mood and neurodegenerative disorders [65]. Furthermore, the analysis method we employed could be useful in studies that test psychological or pharmacological interventions designed to improve well-being [66], allowing these investigations to disentangle whether their intervention targets the underlying neural architecture, changes in patterns of thought, or a combination of both.

## METHODS

### Participant Demographics

In accordance with the Declaration of Helsinki on the conduct of research involving human participants, ethical approval was obtained for this study from the Department of Psychology and York Neuroimaging Centre, University of York ethics committees. Following a standard informed consent procedure, a total of 226 healthy, right-handed (one left-handed), native English speaker undergraduate or post-graduate students with normal to corrected vision were recruited from the University of York. All volunteers received monetary compensation or course credit for their participation in line with the departmental policies. As per the exclusion criteria, none of the participants had a history of psychiatric or neurological illness, severe claustrophobia, anticipated pregnancy or drug use that could alter cognitive functioning. Moreover, an extensive motion correction procedure was followed (described in detail below) that resulted in the exclusion of 12 participants due to excessive head motion inside the scanner, and three participants were removed due to the impartial completion of the thought sampling method. In total, 211 participants imaging and thought sampling data were used in this analysis. The average age for this group was 20.85 years (range = 18 - 31, SD = 2.44) with a 129/82 female to male ratio.

### Decomposition of Thought Patterns

To ascertain the principal dimensions of variation in the thought patterns of this participant cohort, a retrospective thought sampling questionnaire was administered immediately after the resting state fMRI scanning. In this session, the participants were asked to subjectively rate their thoughts during the resting state scan on a 4-choice Likert scale from “Not at all” to “Completely” based on a randomly presented set of questions that probed the content and form of thoughts. This set of questions and the accompanying analysis techniques have been extensively utilized in various thought sampling reports previously published in the literature (Supplementary Table S1) [18, 30, 33, 39, 42]. First, the ratings from each participant were hierarchically clustered based on the similarity of responses using the Ward linkage method (Supplementary Fig. S1a). This technique was utilized in order to partition the thought ratings into two distinct groups, thus reducing the number of variables to be decomposed into interpretable patterns of thought [67]. Subsequently, both groups of ratings were reduced to three factors each (six in total) using principal component analysis (PCA) in SPSS (Version 23) (https://www.ibm.com/products/spss-statistics). The number of components was chosen based on scree-plots indicating the eigenvalue of each subsequent decomposition, and its ability to explain variability in the data (Supplementary Fig. 1b). The component loadings for the total number of six decompositions were then rotated using the Varimax method and the resulting factors were visualized on heat maps (Supplementary Fig. 2a-b) and word clouds (Fig. 2). The component scores of each participant on these patterns of thought were then used as between-subject covariates of interest in the subsequent analyses.

### Well-being Assessment

For the assessment of the participants self-perceived well-being, we employed the brief version of a health questionnaire previously established by the World Health Organization Quality of Life (WHOQOL) group [40]. Termed WHOQOL-BREF, this quality of life assessment has been developed to provide a more comprehensive index of overall health of nations that extends beyond measures of mortality and morbidity, hence, it was designed to be readily administered across cultures and countries with different economic status. Extensively validated in a large number of centres, WHOQOL-BREF consists of 26 questions that broadly group into four domains of physical, psychological, social and environmental health. All participants were asked to complete this questionnaire at a separate session administered outside the scanner. Out of 211 participants, 169 fully completed the questionnaire. The standardized responses for the psychological and social health domains were then used as covariates of interest in subsequent linear regression and mediation analyses.

### MRI Data Acquisition

All MRI data acquisition was carried out at the York Neuroimaging Centre, York with a 3T GE HDx Excite Magnetic Resonance Imaging (MRI) scanner using an eight-channel phased array head coil. A single run of 9-minute resting state fMRI scan was carried out using single-shot 2D gradient-echo-planar imaging. The parameters for this sequence was as follows: TR = 3 s, TE = minimum full, flip angle = 90, matrix size = 64 x 64, 60 slices, voxel size = 3 x 3 x 3 mm3, 180 volumes. During resting state scanning, the participants were asked to focus on a fixation cross in the middle of the screen. Subsequently, a T1-weighted structural scan with 3D fast spoiled gradient echo was acquired (TR = 7.8 s, TE = minimum full, flip angle= 20, matrix size = 256 x 256, 176 slices, voxel size = 1.13 x 1.13 x 1 mm3),

### MRI Data Preprocessing

All preprocessing and denoising steps for the MRI data were carried out using the SPM software package (Version 12.0) (http://www.fil.ion.ucl.ac.uk/spm/) and Conn functional connectivity toolbox (Version 17.f) (https://www.nitrc.org/projects/conn) [68], based on the MATLAB platform (Version 16.a) (https://uk.mathworks.com/products/matlab.html). The first three functional volumes were removed in order to achieve steady state magnetization. The remaining data was first corrected for motion using six degrees of freedom (x, y, z translations and rotations), and adjusted for differences in slice-time. Subsequently, the high-resolution structural images were co-registered to the mean functional image via rigid-body transformation, segmented into grey/white matter and cerebrospinal fluid probability maps, and were spatially normalized to Montreal Neurological Institute (MNI) space alongside with all functional volumes using the segmented images and a priori templates. This indirect procedure utilizes the unified segmentationnormalization framework, which combines tissue segmentation, bias correction, and spatial normalization in a single unified model [69]. No smoothing was employed, complying with recent reports on the negative influence of this procedure on the construction of functional connectomes and graph theoretical analyses [70].

Furthermore, a growing body of literature indicates the potential impact of volunteer head motion inside the scanner on the subsequent estimates of functional connectivity and network neuroscience metrics [71–74]. In order to ensure that motion and other artefacts did not confound our data, we have employed an extensive motion-correction procedure and denoising steps, comparable to those reported in the literature [75, 76]. In addition to the removal of six realignment parameters and their second-order derivatives using the general linear model (GLM) [77], a linear detrending term was applied as well as the CompCor method that removed five principle components of the signal from white matter (WM) and cerebrospinal fluid (CSF) [78]. Moreover, the volumes affected by motion were identified and scrubbed based on the conservative settings of motion greater than 0.5 mm and global signal changes larger than z = 3. A total of 12 participants, who had more than 15% of their data affected by motion was excluded from this study [76]. Though recent reports suggest the ability of global signal regression to account for head motion, it is also known to introduce spurious anti-correlations, and was thus not utilized in our analysis [62, 79, 80]. Nevertheless, the composite motion score (i.e. percentage of invalid scans) for each participant was also added as a covariate in group-level analyses to further account for the potential influence of head motion on functional connectome estimations. Finally, a band-pass filter between 0.009 Hz and 0.08 Hz was employed in order to focus on low frequency fluctuations [81, 82]. The maximum, and mean motion parameters and global signal change, the percentage invalid volumes that were scrubbed, and the distribution of correlation coefficients before and after denoising steps are provided in Supplementary Figure S4.

### Functional Connectome Analysis

#### Brain Parcellation

We adopted a set of 264 regions based on the Power et al. 2011 brain parcellation scheme that has been previously shown to produce reliable network topologies at rest and task conditions [44, 83, 84]. The network partitions outlined by Cole et al. 83 were utilized to pre-assign each one of the 264 ROIs to one of the 13 LSNs documented in the original publication [44]. Namely, 10 well-established networks covering dorsal (DAN) and ventral attention (VAN), salience (SAN), cingulo-opercular (CON), fronto-parietal control (FPN), default mode (DMN), visual (VN), auditory (AN), somatomotor (hand and mouth) (SMN), subcortical networks (SCN); as well as 3 networks that fall into memory retrieval, cerebellum, and a network of uncertain function were used as the 13 network partitions [44].

#### Connectome Construction

Fully-connected, undirected and weighted matrices (264 ROI x 264 ROI) of bivariate correlation coefficients (Pearson r) were constructed for each participant using the average BOLD signal time series obtained from the 6 mm (radius) spheres placed on the MNI coordinates of all the 264 ROIs described above. The matrices reflected both positive and negative weighted correlations. The arbitrary thresholding and binarization processes in graph theoretical analysis often lead to loss of information, especially in the case of negative correlations [85]. Given recent reports suggesting a neurophysiological basis and potential cognitive importance of such anti-correlations in healthy brain function [48, 49, 86], we focused on fully connected, weighted connectomes.

#### Network-Based Statistics

Next, we aimed to ascertain components of individual functional connectomes that significantly predicted the participants between-subject variation on the identified patterns of thought. For that purpose, we employed the Network-Based Statistics (NBS) toolbox (Version 1.2) (https://www.nitrc.org/projects/nbs/) [45], which provides enhanced power to identify connected brain components formed by suprathreshold edge links that are associated with a covariate of interest, while controlling for Family-Wise Error (FWE) at the component-level. Utilizing this method, we entered the individual component scores for all six patterns of thought as variables of interest, while accounting for the effects of mean connectivity, age, gender and percentage of motion-related invalid scans identified by the scrubbing procedure. Using these regressors, T-tests were first carried out on fully connected whole-brain network edges for each pattern of thought to assess the relationship between the strength of an edge link and component scores on patterns of thought, storing the size of the connected components that survived the chosen T threshold. Next, over a total of 5,000 permutations in which the outcome measures were randomised, random null distributions of maximal component size above the chosen threshold was generated. The number of permutations in which the maximal component size was greater than the empirical component size, normalised by the total number of permutations, was used to estimate p-values (.05 level of significance). While the initial T_threshold_ = 3.2 was used for the main analysis, comparable results for T_threshold_ = 3.1 and T_threshold_ = 3.3 are reported in the Supplementary Results section (Supplementary Fig. S5). The resulting connected brain components, the links of which showed a significant relationship with individual variability in thought patterns, were then defined as mask graphs to threshold individual functional connectomes, which were then carried forward onto graph theoretical analyses [87].

#### Network Neuroscience Analysis

Graph theoretical metrics in this study were calculated using MATLAB functions obtained from the publicly available Brain Connectivity Toolbox (https://sites.google.com/site/bctnet/). Commonly used in the identification of hub regions that greatly influence the efficiency of a network in distributing information [88], network strength denotes one of the most fundamental measures in weighted functional connectomes and thus formed the basis of our network neuroscience analysis approach. Calculated as the sum of all neighbouring link weights [88], we measured the strength of connected components across all participants that were previously identified as illustrating significant relations to individual variability in thought patterns. Based on recent reports suggesting the importance of anti-correlations, we calculated positive, negative as well as total strength for each individual. Furthermore, given recent evidence suggesting a contribution of the balance between both positive and negative correlations to healthy brain processing [48, 49, 86], we aimed to utilize a metric that incorporated the importance of the interplay between these links when assessing its predictive power for explaining individual variability in self-reported mental well-being. Thus, we defined fractional strength as the ratio of the sum of positive to negative links, which was later used as the graph metric of interest in subsequent linear regression and mediation analyses. Finally, to identify central nodes in the functionally connected components that significantly related to the thought structures, betweenness centrality, i.e. the fraction of all shortest paths in the network that pass through a given node, was calculated. The average positive and negative links and the employed graph theoretical metrics were visualized on circular plots using Circos [89].

#### Test-Retest Reliability Analysis

Our next aim in this analysis was to determine the reliability of the identified patterns of thought and brain connectivity measures. This would not only establish the generalizability of our results but would also allude to the potential differences in the state versus trait-level variability of our neurocognitive measures. For that purpose, a second session of resting state scan and experience sampling was carried out for 44 participants using the same parameters outlined above. This behavioural and imaging data were preprocessed using the same procedures, resulting in the exclusion of four participants due to excessive motion. For the thought decomposition scores, the hierarchical clustering and PCA decompositions obtained from the initial cohort was imposed on the ratings obtained from the second session [90]. Intraclass correlation coefficients (ICC) were employed to assess the reliability of thought component scores and the associated brain connectivity (as denoted by fractional strength) from the first and second sessions. Furthermore, we assessed a potential link between the change in thought patterns between the two sessions and the change in the associated brain functional connections using Pearson correlations.

#### Mediation Analysis

Finally, we used linear regressions to identify the relationship between component scores on the identified patterns of thought, the fractional strength (natural log) of connected components and the participants self-reported scores on the psychological and social domains of the WHOQOL-BREF questionnaire. The relationships with health were Bonferroni corrected for multiple comparisons across the two health domains. After establishing linear relationships between all three measures, subsequent mediation analyses were carried out with the aim of determining the indirect effect of brain connectivity on psychological and social well-being through participants thought patterns. As it is suggested for small to medium sample sizes [91], the percentile bootstrap estimation approach with 5,000 samples were used to ascertain the presence of a mediation effect.

## ACKNOWLEDGMENTS

This study was supported by the European Research Council (Project ID: 646927) (https://cordis.europa.eu/project/rcn/198057/factsheet/en) awarded to JS and a grant from the John Templeton Foundation, “Prospective Psychology Stage 2: A Research Competition”. The funders had no role in study design, data collection and analysis, decision to publish, or preparation of the manuscript. The authors extend their gratitude to Mladen Sormaz, Charlotte Murphy, Hao-Ting Wang and Giulia Poerio for their invaluable contribution to the scanning of participants. In addition, the authors thank Andre Gouws, Ross Devlin, Jane Hazell and the rest of the York Neuroimaging Centre staff for their support in setting up the imaging protocol and scanning. Finally, we thank all the participants for their time and effort in taking part in this study. The authors declare no conflict of interest.

## Competing Interests

The authors declare no competing interests.

## Author Contributions

D.M., E.J. and J.S. designed the experiment. D.V. and T.K. developed experimental materials, performed data collection and analyzed the data. D.V., T.K., E.J., and J.S. interpreted results. D.V. wrote the manuscript with comments from all other authors.

