## Supplementary Information for "Distinct patterns of thought mediate the link between brain functional connectome and psychological well-being"

### Decomposition of Patterns of Thoughts

The hierarchical clustering of ratings on the 25 questions from the thought sampling questionnaire resulted in two major clusters (Fig. S1a). The two clusters of ratings were then decomposed into patterns of thought using PCA. The eigenvalues scores per component number for both clusters of ratings are provided in Figure S1b. The cut-off point of three components was selected based on the eigenvalue (>1) and the additional explanatory power gained by each additional component. The heat maps for the component loadings of the six identified patterns of thought are given in Figure S2a-b, while Figure S3 displays typical responses from participants who scored highest on a given thought pattern. The component scores for each individual on these patterns were then carried forward on to the NBS analysis.

### MRI Data Quality Assessment

The distributions of maximum and average motion parameter values, as well as the average correlation coefficients before and after the employed denoising procedures are provided in Supplementary Figure S4. Following a strict motion-correction procedure, 12 participants who had more than 15% of their data affected by motion were excluded from the analysis.

### Network-Based Statistics

For the main NBS analysis, t-tests were carried out on fully connected whole-brain networks for each pattern of thought at an initial T threshold of T = 3.2 over 5,000 permutations and $p < .05$ level of significance. For the two patterns of thought which significantly related to brain connectivity components, we also provide two further analyses using T thresholds of T = 3.1 and T = 3.3. This yielded significant and comparable results to the T = 3.2 threshold reported in the main manuscript (Supplementary Fig. S5).

*

TABLE S1. Set of thought sampling questions administered immediately following the resting state scanning session. Participants characterized their thoughts based on a 4-point Likert scale.

| Number | Question | Naming |
|---|---|---|
| 1 | My thoughts involved future events. | Future |
| 2 | My thoughts involved past events. | Past |
| 3 | My thoughts involved myself. | Self |
| 4 | My thoughts involved other people. | Other |
| 5 | I thought about something positive. | Positive |
| 6 | I thought about something negative. | Negative |
| 7 | My thoughts were in the form of images. | Images |
| 8 | My thoughts were in the form of words. | Words |
| 9 | My thoughts were detailed and specific. | Specific |
| 10 | My thoughts tended to evolve in a series of steps. | Evolving |
| 11 | My thoughts were vivid as if I was there. | Vivid |
| 12 | My thoughts were spontaneous. | Spontaneous |
| 13 | My thoughts were deliberate. | Deliberate |
| 14 | This thought was similar to thoughts I often have. | Habitual |
| 15 | My thoughts were related to the here and now. | Here-Now |
| 16 | My thoughts were related to a more distant time. | Distant-Time |
| 17 | My thoughts were hard for me to stop. | Hard to Stop |
| 18 | My thoughts were on topics that I care about. | Important |
| 19 | My thoughts were about ideas rather than events or objects. | Abstract |
| 20 | My thoughts at different points in time were all on the same theme. | Thematic |
| 21 | My thoughts dragged my attention away from the external world. | Decoupling |
| 22 | My thoughts were intrusive. | Intrusive |
| 23 | I was thinking about an event that has happened or could take place. | Realistic |
| 24 | My thoughts gave me a new insight into something I have thought about before. | Insightful |
| 25 | I was thinking about solutions to problems (or goals). | Problem-based |

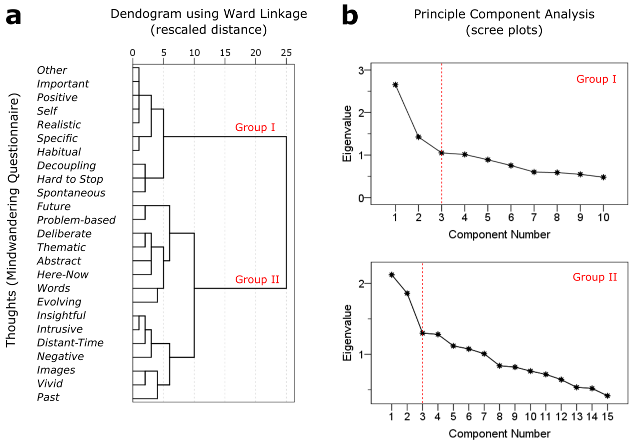


FIG. S1. **Hierarchical clustering and principal component analysis of ratings on the thought sampling questionnaire.** The participants ratings for each question was hierarchically clustered using the Ward linkage method. (*a*) The dendrogram for the resulting clusters indicates two major clusters separated into Group I and II. (*b*) Subsequently, the PCA analysis indicated three components for each group as the optimal number of components with eigenvalue >1, and added explanatory value gained by including an additional component.

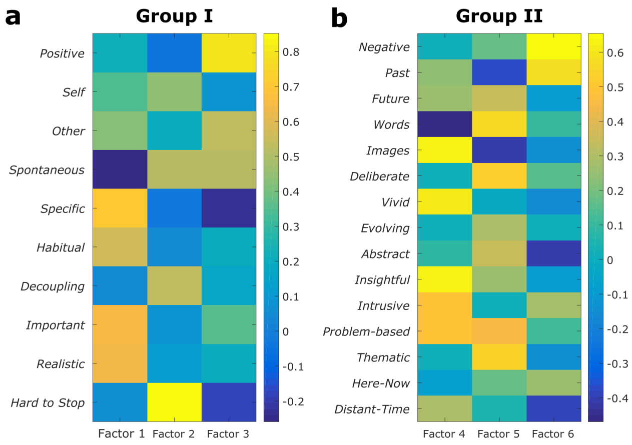


FIG. S2. **Decomposition of patterns of thought.** Two main groups of hierarchically clustered responses to the experience sampling questionnaire were decomposed into three patterns of thought each ($a$-$b$), resulting in a total of six decompositions. Heat maps denote the Varimax rotated component loadings for each decomposition. The component scores for each participant on these patterns of thought where utilized as covariates of interest in the subsequent neuroimaging data analyses.

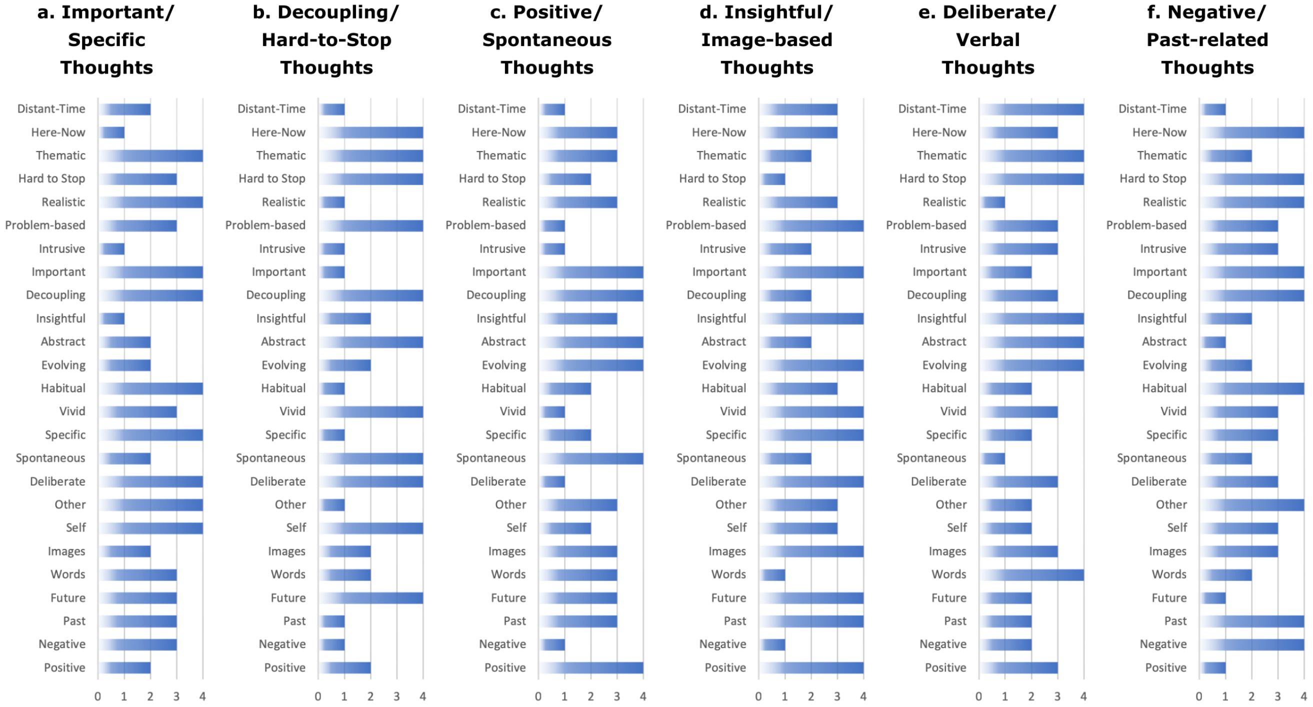


FIG. S3. **Typical ratings on the thought sampling questionnaire.** The bar charts illustrate raw ratings of the participants with the highest component score in the identified patterns of thought ($a$-$f$).

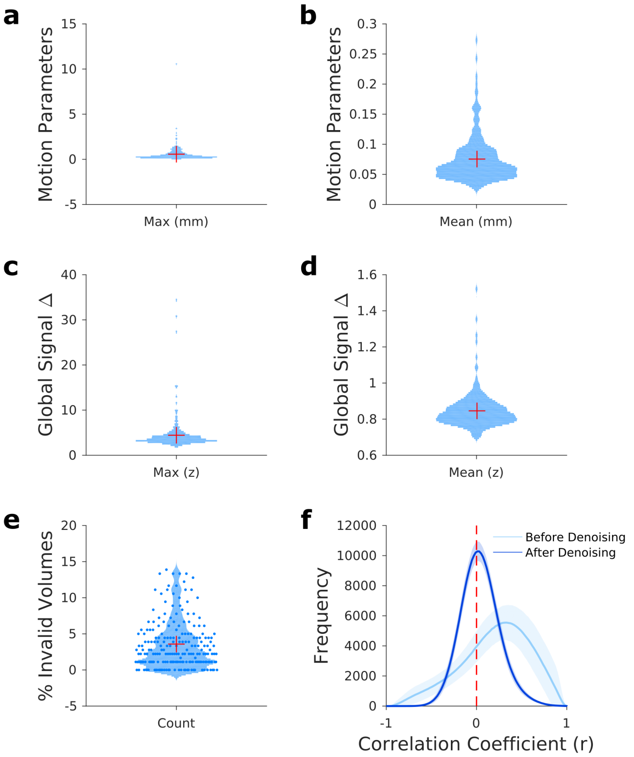


FIG. S4. **MRI data quality assessment and motion correction.** An extensive motion-correction procedure was employed including the removal of motion parameters and their second-order derivatives, CompCor components attributable to white matter and cerebrospinal fluid and linear detrending. In addition, the volumes associated with excessive motion were identified and scrubbed. Participants with a percentage of invalid volumes greater than 15% of their total data were excluded from the analysis. Distributions of (*a*-*b*) mean and maximum translation parameters (mm), (*c*-*d*) mean and maximum global signal change (z), and the (*e*) percentage of invalid scans for the final cohort of participants that were included in this analysis are provided using violin plots. The red stars indicate the 50th percentile. (*f*) In addition, the histogram of the average voxel-based correlation coefficients (r) across participants showed a normal distribution following the denoising steps employed in this study. The shaded areas represent standard deviation.

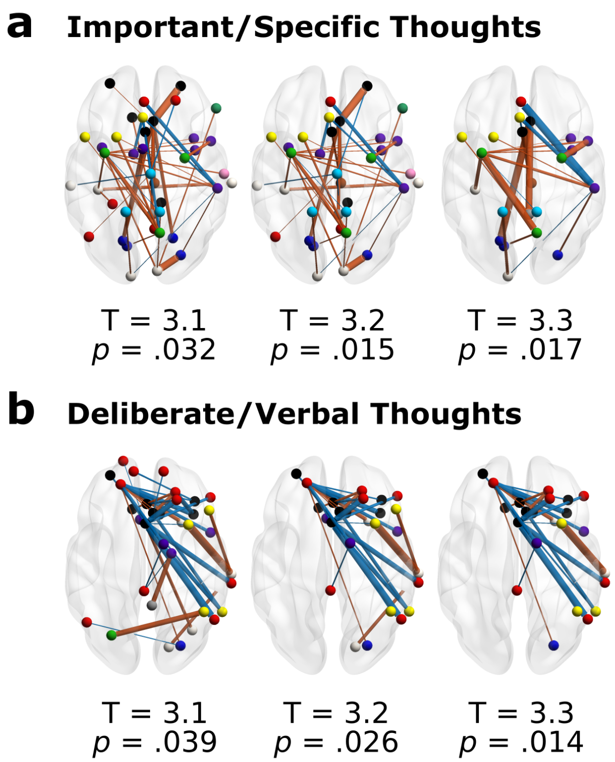


FIG. S5. **Replicability of network-based statistics results at different initial T thresholds.** For the two patterns of thought that significantly related to brain connectivity components, the same NBS analysis was run using two different T thresholds at T = 3.1 and T = 3.3. The resulting brain components are visualized on MNI152 smoothed brains and the corresponding p values of the statistical analysis for the two components (*a*-*b*) and at different thresholds are provided.
